# Discovery of a new song mode in *Drosophila* reveals hidden structure in the sensory and neural drivers of behavior

**DOI:** 10.1101/221044

**Authors:** Jan Clemens, Philip Coen, Frederic A. Roemschied, Talmo Pereira, David Mazumder, Diego Pacheco, Mala Murthy

## Abstract

Deciphering how brains generate behavior depends critically on an accurate description of behavior. If distinct behaviors are lumped together, separate modes of brain activity can be wrongly attributed to the same behavior. Alternatively, if a single behavior is split into two, the same neural activity can appear to produce different behaviors [1]. Here, we address this issue in the context of acoustic communication in *Drosophila*. During courtship, males utilize wing vibration to generate time-varying songs, and females evaluate songs to inform mating decisions [2-4]. *Drosophila melanogaster* song was thought for 50 years to consist of only two modes, sine and pulse, but using new unsupervised classification methods on large datasets of song recordings, we now establish the existence of at least three song modes: two distinct, evolutionary conserved pulse types, along with a single sine mode. We show how this seemingly subtle distinction profoundly affects our interpretation of the mechanisms underlying song production, perception and evolution. Specifically, we show that sensory feedback from the female influences the probability of producing each song mode and that male song mode choice affects female responses and contributes to modulating his song amplitude with distance [5]. At the neural level, we demonstrate how the activity of three separate neuron types within the fly’s song pathway differentially affect the probability of producing each song mode. Our results highlight the importance of carefully segmenting behavior to accurately map the underlying sensory, neural, and genetic mechanisms.

## Results

### Drosophila melanogaster courtship song comprises at least three, not two, distinct song modes

Manual inspection of song recordings have identified two distinct song modes in *Drosophila melanogaster:* sine song, which consists of a sustained sinusoidal oscillation, and pulse song, which comprises trains of short impulses (ref). Based on this observation, automated methods were developed to effectively detect and segment these two modes of song (Arthur et al.), and using this software, large data sets have been analyzed to link both genes and neural activity with song patterns [5-9]. However, the claim that song comprises only two modes has never been tested using statistical methods, leaving open the possibility that more than two modes exist (as has been suggested for other *Drosophila* species, [10]). We therefore took a more unbiased approach and clustered 25 millisecond waveforms of previously collected raw song recordings [9]. We first classified each sample as signal or noise based on signal amplitudes that exceeded background noise (33% of the waveforms passed this criterion); no other constraints on waveform shape were imposed (Figure 1A, see STAR Methods for details). We then aligned the signal samples to their peak energy, normalized them to correct for differences in waveform amplitude (induced by variations in the intensity of male singing or his position relative to the microphone), and adjusted their sign so that the waveform was positive immediately preceding the peak (this was done because waveform inversions are likely caused by changes in male position relative to the microphone) (Figure 1B). The resulting set of ∼20,000 normalized signals from 47 wild type males of the strain NM91 contained courtship song as well as non-song noises originating from grooming, jumping or other behaviors.

**Figure 1:**
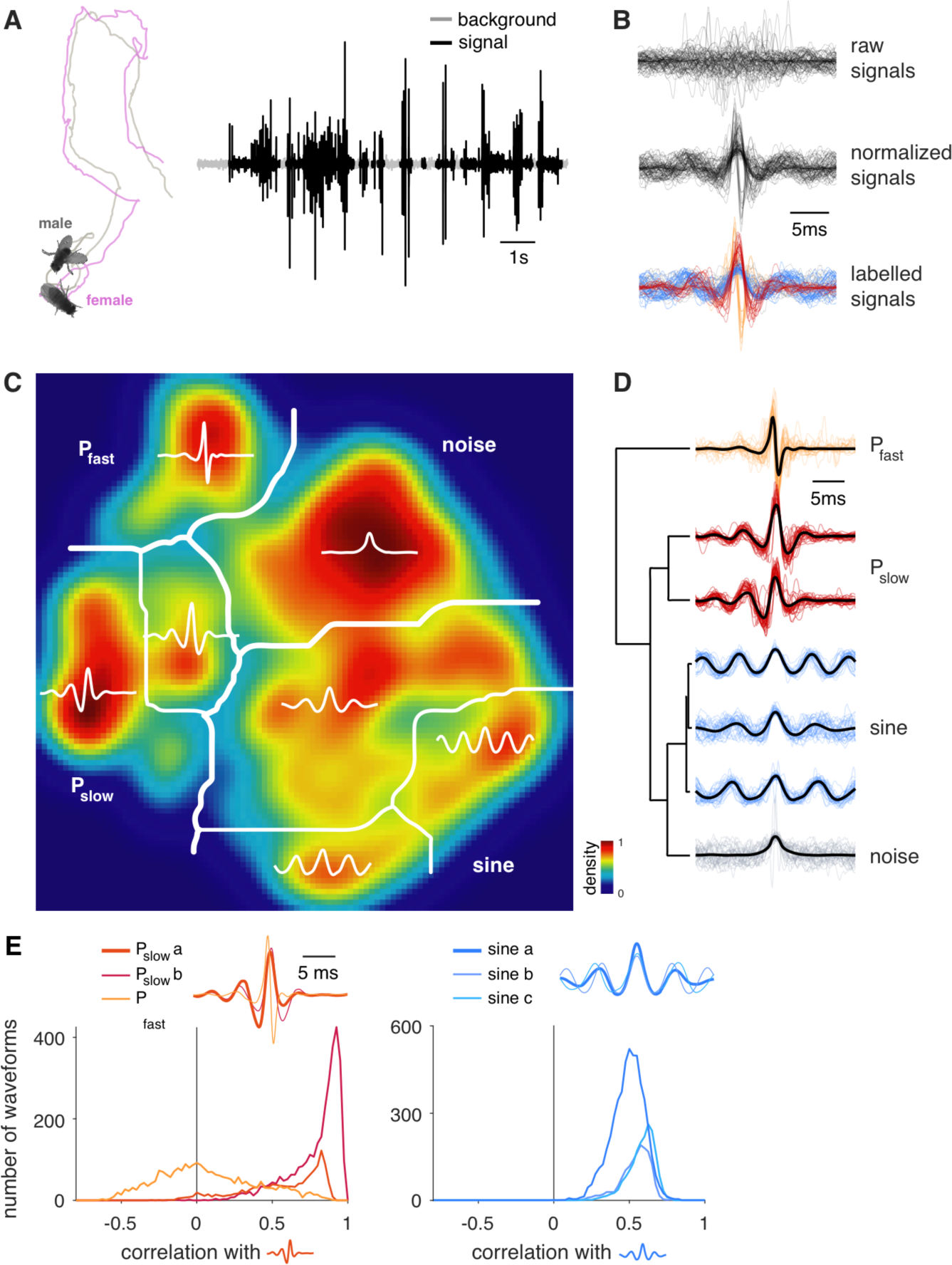
*Drosophila melanogaster* courtship song comprises three, not two, modes. **A** During courtship, the male chases the female and produces a courtship song through unilateral wing vibrations. Shown are male (gray) and female (magenta) traces during courtship. Song recording (right) with background noise (grey) and signals (black) from a wild type *D. melanogaster* male. **B** Non-overlapping signal chunks (25ms duration) were extracted (top) and then normalized (middle) to scale all signals to the same peak amplitude and to flip the sign of signal waveforms such that the largest peak prior to the pulse center is always positive. Variation in signal amplitude or sign could be due to the male’s position relative to the microphone. Here we show a random subset of 86 out of all 21,104 chunks. After clustering, signals produced by males during courtship fall into four classes: non-song noises (grey), sine (blue), P_slow_ (red) and P_fast_ (orange). **C** We reduced the dimensionality of 21,104 waveforms from 47 individual males of the *D. melanogaster* strain NM91 using t-distributed stochastic neighbor embedding (tSNE) and the resulting two-dimensional distribution (density color coded, see colorbar) of signals was partitioned using the watershed algorithm. This yielded seven clusters, which upon inspection of the waveforms, correspond to noise, sine song (three clusters) and two distinct pulse song modes - P_slow_ (two clusters) and P_fast_ (see text labels). Thick white lines mark the main mode boundaries after cluster consolidation (see D) and thin white lines mark the submodes within each main mode. **D** Hierarchical clustering of the average waveform for each watershed cluster (centroids, thick black lines) supports the grouping of clusters into 4 main modes. Horizontal distance in the tree (left) corresponds to the dissimilarity between cluster centroids. Thin colored lines (right) show individual waveforms for each cluster. The waveforms corresponding to non-song noises (grey) are highly heterogeneous and have in common only the peak in energy to which they were aligned. Three sine song clusters (blue) mainly differ in frequency and constitute a continuum of waveforms. Note the second sine cluster is heterogeneous. Two P_slow_ clusters (red) differ only in their asymmetry, with more weight on either the negative lobe leading (bottom) or lagging (top) relative to the main positive peak. P_fast_ (orange) joins all fast and biphasic pulses. **E** Similarity between all pulse waveforms (left) or all sine waveforms (right) obtained from the watershed clustering (Figure 1C). Similarity was computed by correlating all pulse or sine waveforms with the centroids for one of the clusters (the waveform used for correlation is thicker in the inset). Distributions do not strongly depend on which of the clusters was chosen as a template for correlation. The distribution for the pulse clusters is bimodal: the two P_slow_ clusters strongly overlap and have high correlation values with the centroid, indicating that they belong to a single song mode. The P_fast_ pulses exhibit projection values onto the P_slow_ centroid symmetrically distributed around 0, indicating very low similarity with the P_slow_ pulses and supporting its classification into a distinct pulse song mode. By contrast, the distribution for all three sine song clusters (right) is skewed towards high values and all three distributions overlap strongly, indicating that sine song constitutes a single song mode.

To facilitate the classification and the visualization of the waveforms, we reduced the dimensionality of the data set using t-distributed stochastic neighbor embedding (tSNE) [11,12]. The tSNE method is a nonlinear dimensionality reduction method that preserves local similarity structure in a data set and is therefore particularly suited for classification. We clustered the low-dimensional representation of our signals in two steps. First, we partitioned the signal distribution along local minima using the watershed algorithm (Figure 1C). This procedure initially yielded seven clusters which represent an over-partitioning of the waveform space, since it cuts the signal space along relatively weak local minima and thereby assigns relatively similar waveforms to different clusters. We therefore employed a second hierarchical clustering step to consolidate the watershed clusters based on the similarity of their centroids (Figure 1D) and we chose the number of modes based on the similarity of the waveforms in the watershed clusters (Figure 1E). This resulted in four distinct signal modes (Figure 1D): i) A “noise” mode lumps all signals that lack common structure across exemplars. ii) A “sine” mode joins three clusters that contain waveforms with sustained oscillations that tile a continuum of carrier frequencies between 120 and 180Hz (Supplemental Figure 1B). The three sine clusters all contain similar waveforms as indicated by the strongly overlapping and high correlation values (Figure 1E, right). This justifies them being merged into a single sine song mode by the hierarchical cluster algorithm. iii) A “pulse” mode with relatively slow (200-250Hz) and symmetrical waveforms connects two very similar clusters with either a stronger leading or lagging lobe. These two clusters form a single pulse mode, since the waveforms are highly correlated (see Figure 1E, left), and they are thus joined by the hierarchical cluster algorithm. iv) A second “pulse” mode that in contrast to the previous one is biphasic (asymmetric) with faster oscillations (250-400Hz). This pulse mode is highly dissimilar with the previous pulse mode (correlation values centered around zero, Figure 1E, right) indicating that it forms a distinct pulse mode. We term the two pulse modes “P_slow_” and “P_fast_”, accordingly. Note that while there is considerable variability within each song mode, the modes are clearly distinct from each other.

To facilitate analyses of the production, perception and evolution of these two pulse types within *Drosophila*, we developed a simpler and more efficient pulse type classifier. The method takes as input pulses and sines detected from an automatic song segmenter [13] - which can detect both pulse types with high speed and reliability (detection rates: 78% for all pulses, Supplemental Figure 1B, 80% for P_slow_, 77% for P_fast_) - and then classifies returned pulses based on their similarity with templates derived from the tSNE analysis pipeline (Figure 2A and Supplemental Figure 1C, see STAR methods for details). This classifier reliably and efficiently reproduces the classification from the full analysis pipeline into the slow, symmetrical P_slow_ and the fast, asymmetrical P_fast_ in all individual males of eight geographically diverse wild type strains, suggesting that the existence of two pulse types is common across isolates of *Drosophila melanogaster* (Figure 2B, Supplemental Figures 1D, 2A, B).

**Figure 2:**
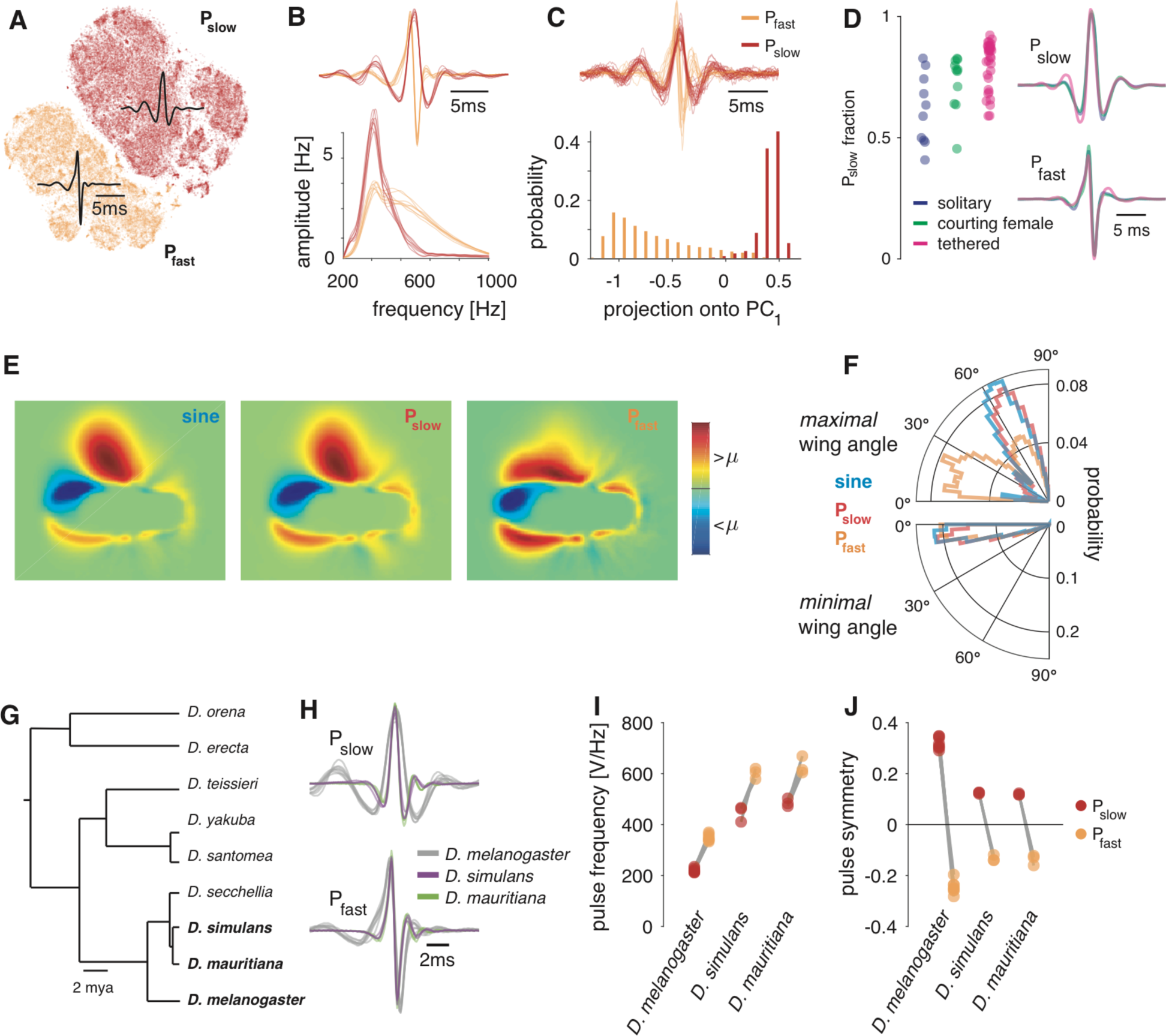
Song pulses can be separated into P_fast_ and P_slow_ across strains and species, and each corresponds to a distinct wing pose. **A** Two-dimensional tSNE of all pulses detected by the automated segmenter reveals a clear separation of the detected pulses into P_slow_ (red) and P_fast_ (orange). Shown are 71,029 pulses from 47 males of the *D. melanogaster* strain NM91. Black waveforms correspond to the average waveform of all pulses classified as P_slow_ (top) or P_fast_ (bottom), respectively. **B** Average waveforms (top) and spectra (bottom) of P_fast_ and P_slow_ for all eight *D. melanogaster* strains investigated (see Methods for strain names). **C** The P_fast_ versus P_slow_ distinction is evident upon inspection of individual pulses (top). Shown are 32 randomly selected, normalized pulses from NM91 males. Both pulse types are also separable using a linear dimensionality reduction technique - PCA (bottom). Plotted is the distribution of projection values of all 71,029 NM91 pulses onto the first principal component. Pulses (top) and projection values (bottom) are colored by the pulse type they were classified as (see legend). **D** Pulses driven by thermogenetic activation of P1 neurons [1] in either solitary males (purple), males courting a female (green), or males tethered and walking on a ball (pink). Flies experienced open-loop visual stimulation matching the natural statistics of female motion on the male retina during courtship [2]. In all three conditions, both pulse types occur though with varying probability (left). Pulse shapes for P_slow_ (top right) and P_fast_ (bottom right) are nearly indistinguishable for all three conditions. **E** Pose of flies during the production of sine, P_slow_ and P_fast_. Shown is the average, mean subtracted frame at the time of the peak amplitude of sine and pulses, respectively (100Hz frame rate, 25 px/mm). Frames were oriented such that the male faces rightward, and flipped such that maximally extended wing is up. Both sine (left) and P_slow_ (middle) are produced with a strongly extended wing while P_fast_ (right) is produced with the wing held closer to the body. **F** Polar histograms of the angle of the maximally (top) and minimally (bottom) extended wing during sine (blue), P_slow_ (red) and P_fast_ (orange) production. Distribution of the maximal wing angles for sine and P_slow_ are similar (50-70∘). The distribution of P_fast_ angles has a main mode at 5-30∘ and a smaller second mode overlapping with the angles observed for sine and P_slow_. Minimal wing angles (bottom) overlap for all three song modes. *Drosophila melanogaster* song is thought to be produced by unilateral wing vibration - the minimally extended wing is therefore likely silent. Data in E, F: A sine, B P_slow_ and C P_fast_ events for from 34 individuals of the *D. melanogaster* strain NM91. **G** Phylogenetic tree of the *melanogaster* species subgroup (reproduced from [3]). The species investigated here are highlighted in bold font. **H** Average pulse waveforms for all strains analyzed, colored by species (see legend). Each strain of each species produces a slower symmetrical (P_slow_, top) and a faster asymmetrical pulse (P_fast_, bottom) type. **I, J** Pulse frequency (I) and symmetry (J) for the pulse waveforms in H by pulse type. *D. simulans* and *D. mauritiana* pulses are faster than those of *D. melanogaster*. For N flies and pulses see Supplemental Figure 2.

We next ran several control analyses to ensure that the two pulse types were not data recording or analysis artifacts. First, we were able to recover both pulse types when using linear (as opposed to nonlinear) dimensionality reduction (principal component analysis instead of tSNE, Figure 2C) and also when omitting parts of the normalization procedure, demonstrating that our finding does not crucially depend on the details of our analysis pipeline (Supplemental Figure 2C). In addition, the existence of two pulse types could be due to the male changing his position relative to the directional microphones used to record song. However, when we induced singing via thermogenetic activation of P1 song pathway neurons in a tethered male walking on a spherical treadmill [5] - thereby fixing his position relative to the microphone - we still observed both pulse types (Figure 2D). Lastly, we examined the fly’s pose during singing and found that the two pulse types are associated with distinct, largely unilateral, wing positions (Figure 2E-F): P_slow_ is produced with one wing fully extended (ca. 70∘) while P_fast_ is produced most often with one wing only weakly extended (ca. 20∘). Together, these analyses demonstrate that the P_slow_ and P_fast_ are separate song modes, likely to be produced by separate motor programs.

What is the function of the minimally extended wing? To answer that question, we recorded song from males that had one wing cut (either left or right). These males showed a reduction of singing to ∼50% of intact males (Supplemental Figure 2D). While these males still produced both pulse types, the individuality of these pulses was strongly reduced (Supplemental Figure 2E, F). This suggests that the existence of two wings increases the individuality of pulse shapes produced by males and hence the function of the 2^nd^ wing may be to make the pulse shape more idiosyncratic. However, we observed no strong effect of cutting one wing on pulse variability (Supplemental Figure 2G).

How do the two pulse types - P_fast_ and P_slow_ - in *Drosophila melanogaster* compare to the pulse types produced in close relatives [10]? We applied our analysis pipeline to song data from two related species, *Drosophila simulans* and *Drosophila mauritiana* (Figure 2G). Since the pulses in *D. simulans* and *D. mauritiana* are about twice as fast as *D. melanogaster*, we adapted the automated song segmentation and pulse normalization procedure to reliably detect and classify pulses in these related species (see STAR Methods for details, Supplemental Figure 1B). Similar to *D. melanogaster*, pulses reproducibly clustered into two major pulse types for all strains of *D. simulans* and *D. mauritiana* examined (Figure 2H). One cluster contains slow, symmetrical pulses and the other faster, asymmetrical pulses (Figure 2H-J, Supplemental Figure 2A, B). *D. melanogaster* pulses are distinguishable from *D. simulans* and *D. mauritiana* pulses only by frequency: P_slow_ and P_fast_ are shifted from ∼220 to ∼470 Hz and from ∼350Hz to 670 Hz with only small differences between *D. simulans* and *D. mauritiana* (Figure 2J). The existence of P_fast_ and P_slow_ in all three species supports the following model for the evolution of pulse shape in the *melanogaster* species group: *D. simulans* and *D. mauritiana* increased the speed with which two ancestral motor primitives - one for each pulse type - were executed. In the sections below, we investigate how the existence of two distinct pulse modes affects our understanding of courtship song patterning and perception in *D. melanogaster*.

### Pulse type choice contributes to the modulation of song amplitude with distance

Males structure their songs into bouts composed of trains of sines and pulses [13], with the choice to sing either sine or pulse biased by sensory feedback from the female [9]. How does classifying song now into three modes affect the patterning of song? We found that males do not continually switch between P_fast_ and P_slow_, but compose pulse trains of one type or the other more frequently than expected by chance (Figure 3A-B). By employing Generalized Linear Models [9,14] to predict pulse type from sensory and movement features (see STAR methods), we found that the distance between the male and female was the strongest predictor of pulse type (Figure 3C). Males from eight different wild type strains consistently bias toward producing P_fast_ at larger distances from the female (Figure 3D). In addition, we observed that blind males (but not deaf or pheromone insensitive males) produce a significantly higher fraction of P_fast_ pulses (Figure 3E), indicating that vision plays a role in the choice between P_slow_ and P_fast_. This builds on previous work that demonstrated males produce sine song when close to the female [9] - we now see that males continually bias toward modes with faster - and higher intensity —wing movements, as their distance from the female increases (Figure 3F). We tested each pulse mode for evidence of amplitude modulation with distance [5] and found that while P_fast_ amplitude increases with distance (males ratchet up P_fast_ amplitude as they get farther from the female without changing pulse shape (Figures 3H,J), P_slow_ amplitude modulation was comparatively weak (Figure 3G). Taken together, these data suggest a reinterpretation of previous work on amplitude modulation [5]: the behavior is driven by at least two processes—the choice to sing the softer or the louder pulse type (P_slow_ vs P_fast_) coupled with analog modulation of P_fast_ amplitude.

**Figure 3:**
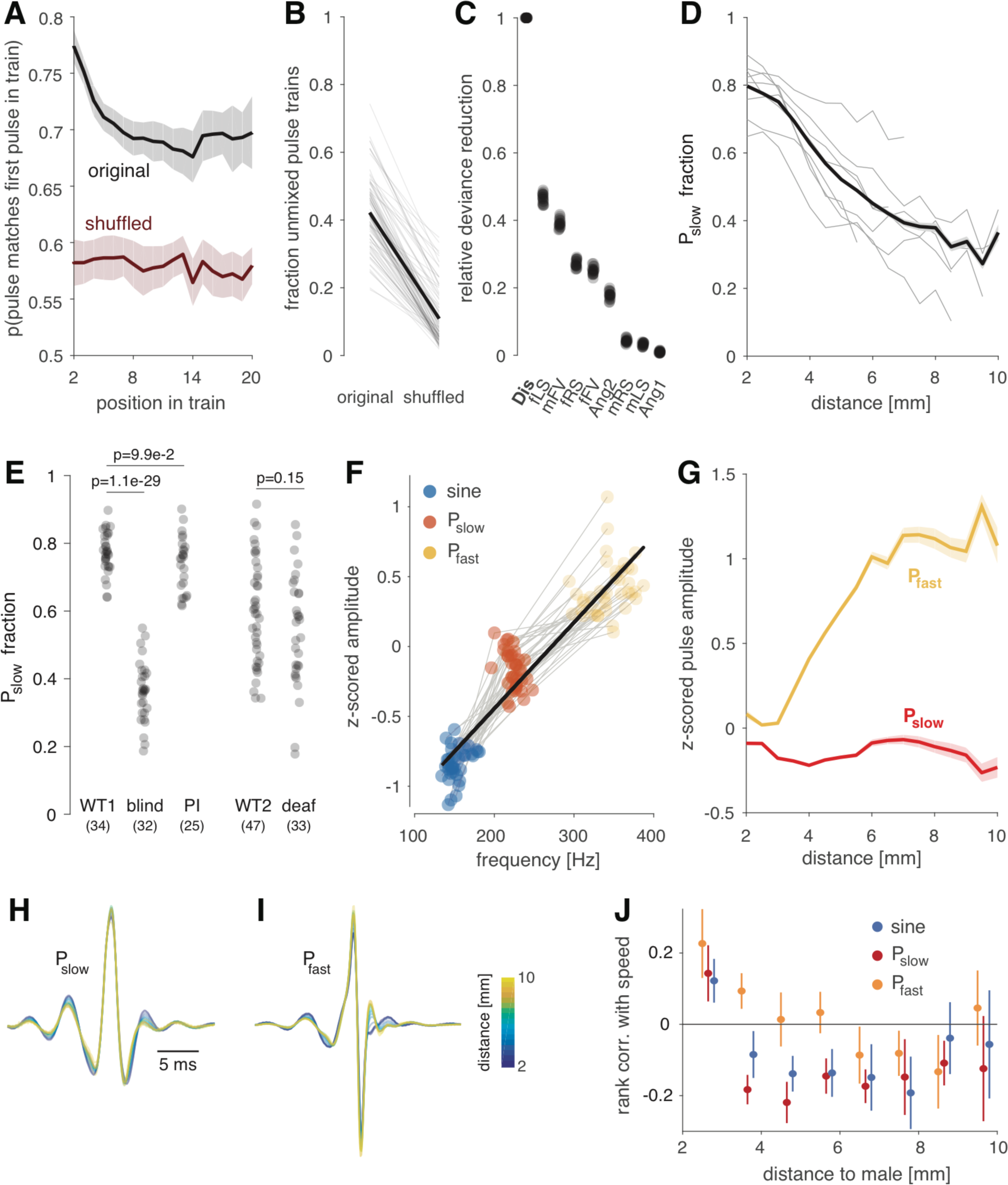
The choice between P_slow_ and P_fast_ is driven by sensory feedback and this choice impacts female responses. **A** When starting a pulse train with one pulse type, males continue to sing that type more than expected by chance. To estimate chance level, the sequence of pulse types was randomly permuted (shuffled) for each fly. Mean ± SEM across 8 *D. melanogaster*strains. N=315 flies. **B** The persistence observed in (A) leads to more pulse trains containing only one pulse type than would be expected by chance (original: 0.42±0.11, shuffled: 0.10±0.07, p=1.7 10^-29^, rank sum test, N=315 flies). Thin lines show the probability, for several individuals, of observing unmixed pulse trains with at least 3 pulses using original (left) and shuffled (right) pulse labels. The thick black line corresponds to the average over all flies. **C** Relative deviance reduction in a generalized linear model that predicts the choice between pulse types (P_slow_ versus P_fast_) from different sensory features (see Methods for details). Distance is most predictive of pulse choice. Each dot corresponds to the performance obtained for a fit to subset (80%) of the data. **D** Fraction of P_slow_ out of all pulses produced as a function of distance. Flies bias towards P_slow_ when close to the female (r^2^=0.92, p=1.0x10^-9^). Thin lines correspond to individual *D. melanogaster* strains, and the thick shaded line depicts mean ± SEM. **E** Fraction of P_slow_ in flies with different sensory manipulations. Blind flies but not pheromone insensitive (PI) or deaf flies produce more P_fast_, indicating that distance estimation requires visual cues. N flies indicated in parentheses and p-values are from a two-tailed t-test. **F** A song mode’s carrier frequency strongly correlates with its amplitude (r^2^=0.86, p=2×10^-51^). Dots correspond to the mean carrier frequency and amplitude (peak to peak) of sine (blue), P_slow_ (red) or P_fast_ (orange) from 47 males of the *D. melanogaster* strain NM91. Grey lines connect song modes of individuals, thick black line is the result of linear regression. **G** P_fast_ (orange) is louder than P_slow_ (red) at all distances and is the only pulse type that is amplitude modulated (P_fast_ r^2^=0.86, p=1.1×10^-7^, P_slow_ r^2^=0.03, p=0.5) (mean ± SEM across 8 *D. melanogaster* strains). **H, I** Average P_slow_ (H) and P_fast_ (I) shape produced at different distances from the female (see colorbar). N= 330759 pulses from 315 individuals and 8 different *D. melanogaster* wild types strains. The shape of both pulse types changes only minimally with distance. **J** Rank correlation between female speed and the amount of sine song (blue) or the number of P_slow_ (red) or P_fast_ (orange) pulses per 30 second time window as a function of distance (mean ± SEM across 8 strains). Sine and P_slow_ are negatively correlated with female speed for almost all distances. P_fast_ only slows females down for distances between 6 and 9 mm. Note the positive correlation when the male is very close to female for all song modes.

### Females slow in response to all three song modes

The correlation between the frequency of a song mode and its amplitude (Figure 3F) suggests that males may switch between song types simply as a strategy to produce song signals of different intensities. The modes, however, could also exert different effects on female behavior. Previous studies have shown that both sine and pulse song are correlated with a reduction of locomotor speed in sexually receptive females, and that this correlation depends on females being able to hear the male song [6,9]. We examined female locomotor speed relative to the amount of each song mode the male produced (per 30 seconds of courtship - previous studies showed that the strongest correlation between male song and female speed was on timescales of tens of seconds [6]). We found that all three song modes were correlated with a reduction in female speed, but in a context-dependent manner. P_slow_ and sine were correlated with reductions in female speed across a wide range of distances, whereas P_fast_ was correlated with a reduction in female speed, but only when males were far away from females (and then only weakly (Figure 3J)). All three modes were correlated with increases in female speed when produced very close to the female. These results only emerge when examining correlations relative to male-female distance (Supplemental Figure 3A-C). That P_slow_ and P_fast_ differentially affect female behavior suggests that they have different “meanings” for the female. Further elucidation of the differential roles of P_slow_ and P_fast_ on female behavior will likely require higher resolution readouts of female postural movement changes [12,15]. The reclassification of male courtship song therefore also has implications for the study of auditory perception in the female nervous system.

### Song pathway neurons modulate the choice between song modes

Above, we showed that males select between pulse types based on sensory feedback (Figure 3C, D), and that these two pulse types correspond to different wing poses (Figure 2E). This suggests that neurons within the fly’s song pathway can bias the motor output towards one of the three song modes. To test this hypothesis, we focused on three previously characterized neurons implicated in driving wing extension or generating pulse song (Figure 4A, Supplemental Figure 4B): P1, a cluster of ∼20 male-specific *Fruitless*+ and *Doublesex+* neurons per hemibrain [16,17], pIP10, a pair of male-specific *Fruitless*+ neurons that descend from the brain to the ventral nerve cord (VNC) [18], and ps1, a *Doublesex+* wing muscle motor neuron in the ventral nerve cord [8].

**Figure 4:**
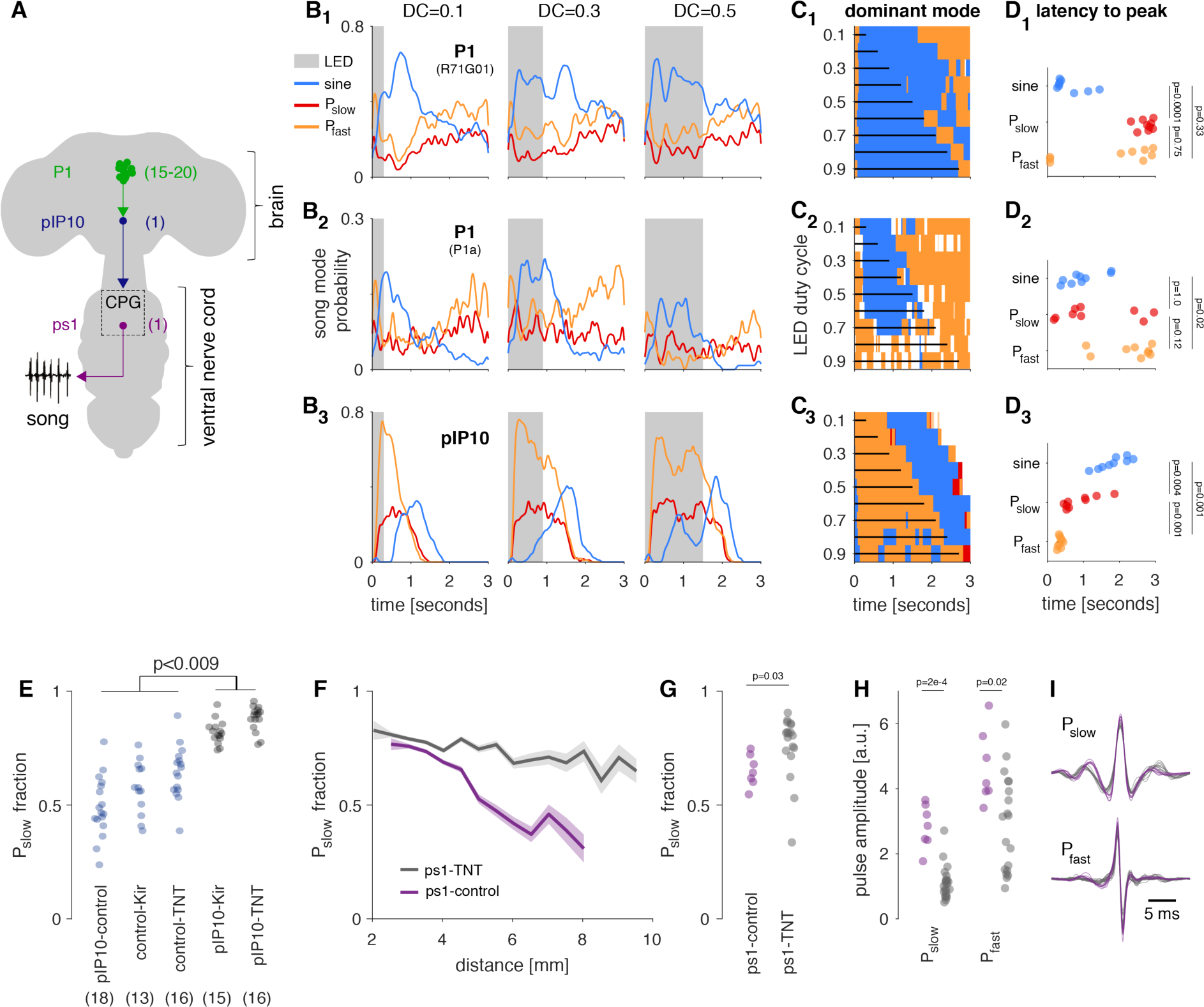
The activity of song command neurons and wing muscles affects the choice between song modes. **A** Schematic of three elements of the putative song pathway [1,4] manipulated in this study. The total number of neurons per hemisphere is indicated in parentheses. P1 neurons are activated by female cues and are local to the brain. We used two separate drivers to label P1 neurons (R71G01 [5] and P1a [6] - see pictogram in Supplementary Figure 4B for more information on which neurons are labeled by these drivers). pIP10 is a descending neuron. ps1 is a motor neuron that innervates the ps1 wing muscle [4]. **B_1-3_** Population average song probability for producing sine song (blue), P_slow_ (red) or P_fast_ (orange) upon optogenetic activation of P1 (R71G01) (B_1_), P1 (P1a) (B_2_), and pIP10 (B**3**). The trials lasted 3 seconds with the LED being activated (shaded gray area) for 300ms (duty cycle (DC) 0.1) and 2700ms (DC 0.9). Shown are data for three out of nine DCs tested (see Supplemental Figure 4A for the full data set). B-D: Average over 6 P1 (R71G01), 6 P1 (P1a), 3 pIP10 flies, 30 trials per fly and DC. **C_1-3_** The dominant song mode (color coded, see legend in A) as a function of time for all nine LED DCs tested. Vertical black lines indicate the duration of the stimulation. When the LED is on, P1 (R71G01) (C_1_) and P1 (P1a) (C__2__) stimulation strongly drives sine song production, while pIP10 (C_3_) drives mainly P_fast_ (orange). Sine song is produced after LED stimulation in pIP10. **D_1-3_** Latency to peak probability for P_slow_ (red) and P_fast_ (orange) for all nine DCs tested (p-values in the plot are the result of two-tailed Wilcoxon rank sum tests between all pairs, corrected for multiple comparisons using the Bonferroni method). pIP10 activation consistently drives P__fast__ with shorter latency than P_slow_ (D_3_). There is no such consistent difference in latency between pulse types for P1 (R71G01) (D_1_) and P1a (D_2_). **E** P_slow_ fraction produced by males courting a female when pIP10 is inactivated (gray, using TNT or Kir2.1, respectively) and in control flies (blue). pIP10 inactivation during courtship leads to significantly more P_slow_ (ANOVA (p=6×10^**-17**^) followed by a Tukey-Kramer post hoc test, p-value is corrected for multiple comparisons). Number of flies for each genotype given in parentheses. **F** Fraction of P_slow_ pulses out of all pulses produced as a function of distance to the female in ps1-control flies (purple, inactive TNT expressed in the ps1 motorneuron) and ps1-inactivated flies (gray, functional TNT expressed in ps1). Shaded lines indicate mean ± SEM across flies). The magnitude of the modulation of pulse choice with distance is strongly reduced upon ps1 inactivation (ps1-control: r^**2**^=0.92, p=3×10^**-5**^; ps1-TNT: r^**2**^=0.72, p=7×10^**-7**^). **G** P_slow_ fraction in ps1-TNT flies (gray) and ps1-control flies (purple). Ps1 inactivation reduces the fraction of P_slow_ pulses produced (p=0.03, two-tailed Wilcoxon rank sum test). **H** Raw microphone amplitude of P_slow_ (left) and P_fast_ (right) pulses in ps1-TNT (gray) and ps1-control (purple) flies. Ps1 inactivation reduces the amplitude of both pulse types (p=2×10^**-4**^ for P_slow_ and 0.02 for P_fast_, two-tailed Wilcoxon rank sum test). **I** Shapes of the P_slow_ (top) and P_fast_ (bottom) pulses in ps1-ctrl (purple) and ps1-TNT flies (gray). Lines correspond to averages over pulses of each type for each fly. P_slow_ slows and widens weakly with ps1 activation. F-I: N=7 control flies (23636 pulses) and 18 experimental flies (14417 pulses).

Either P1 or pIP10 neural activation results in song production, even in the absence of a female [5,18]. P1 neurons are known to be involved not only in song production, but also in the integration of pheromonal and visual signals from the female [17,19-21], the production of aggressive behaviors [22], and overall mating drive [23]. We reasoned that when males are close to females, P1 activity should be elevated, since the pheromonal and visual cues provided by the female should be particularly strong. Since, males bias towards both sine song and P_slow_ production when close to the female ([9], Figure 3D), optogenetically activating P1 neurons should preferentially drive either sine or P_slow_, but not P_fast_, even in the absence of a female. pIP10 neurons are thought to be postsynaptic to P1 neurons [18], and therefore might exert a similar effect on song patterning.

To investigate how the activation of P1 and pIP10 affects the probability of producing the three different song modes, we used csChrimson [24] to optogenetically stimulate each neuron type in solitary males. We chose an LED stimulation protocol in which we tested varying duty cycles (DCs) between 0.1 and 0.9 during three-second long trials, thereby leaving either very long pauses (DC=0.1 corresponds to a pause of 2700ms) or very short pauses (DC=0.9 corresponds to a pause of 300ms) between subsequent stimulations - in other words, short DCs correspond to weaker stimulation and long DCs to stronger stimulation. Consistent with the idea that P1 encodes female proximity, males produced mainly sine song during P1 activation, while both P_fast_ and P_slow_ were largely suppressed (Figure 4B_1,2_, 4C_1,2_). We observed this effect for two different genetic drivers that target P1 neurons: R71G01 [25] and a split GAL4 driver termed P1a [22] (Supplemental Figure 4B). Interestingly, for P1 a, sine song was only dominant for DCs <0.8; at higher DCs, pulse song was more dominant, suggesting sine production requires a pause between stimulations (Supplemental Figure 4A2, Figure 4C_2_). P_fast_ and P_slow_ probabilities exhibited similar dynamics for both P1 drivers (Figure 4B_1,2_, Supplemental Figure 4A_1,2_) suggesting that P1 mainly biases the choice between sine and pulse song, while affecting the balance between P_fast_ and P_slow_ relatively little. Moreover, P1 activation did not affect the waveform shape of either P_fast_ and P_slow_, only the probability of producing pulses (Supplemental Figure 4C, D).

pIP10 is a descending neuron thought to be postsynaptic to P1 [18]. However, we found that pIP10 does not simply relay P1 activity, since P_fast_, and not sine song, was the dominant mode produced during optogenetic activation (Figure 4B_3_, 4C_3_). Sine became the dominant song mode only after stimulation ended and only if the pauses between stimulation were sufficiently long (DCs ≤ 0.9). Consistent with these results, we also found that pIP10 inactivation (using either Kir2.1 [26] or TNT [27]) during courtship with a female decreased P_fast_ production (Figure 4E). We found that the probability of producing P_slow_ was largely independent of optogenetic stimulus duration, whereas the opposite was true for P_fast_: for short DCs (<0.6) we observed high peak probabilities, intermediate DCs (0.6-0.8) produced an onset transient with reduced steady-state levels, and at the longest DCs, P_fast_ levels were relatively constant throughout the stimulation period and resembled those of P_slow_ (Figure 4B_3_, Supplemental Figure 4A_3_).

Our results with P1 and pIP10 activation highlight the necessity of carefully delineating behavioral modes and reveal complex dynamics in the neural circuits driving song production. For activation of both cell types, for example, we observed that stimulus history (and thereby the history of song produced) has a strong impact on the song types elicited by optogenetic activation (Figure 4B, Supplemental Figure 4A). While optogenetic activation of both P1 and pIP10 affected male speed (Supplemental Figure 4A), we observed no fixed relationship between male speed and song choice among the three drivers examined here, suggesting that the song dynamics induced by optogenetic activation cannot be explained as an indirect effect of changes in male speed alone.

How does the activity of P1 and pIP10 neurons ultimately lead to changes in the choice of pulse types? While much of the song pathway has not yet been mapped [28], two sets of motor neurons have been shown to be involved in either sine or pulse song [8]. Inactivation of the ps1 motor neuron reduces both pulse amplitude and carrier frequency [8]. This effect would be consistent with ps1 being involved in P_fast_ production, since P_fast_ is the louder and faster of the two pulse types. Since activation of ps1 alone does not generate song, we instead silenced ps1 using tetanus toxin (TNT), and paired males with females to determine the effect of ps1 silencing on song production. We found that ps1 inactivation does not have a strong effect on the shape of either P_slow_ or P_fast_ (Figure 4H), and only a subtle effect on the frequency of P_slow_, but not P_fast_, pulses (Supplemental Figure 4E, F). Instead, silencing ps1 strongly affects the switching between P_slow_ and P_fast_ with distance (Figure 4F), and thereby reduces the amount of Pf_a_st pulses being produced (Figure 4G): wild type flies bias away from P_slow_ with increasing distance but ps1 inactivated flies continue to sing P_slow_ even when far from the female. ps1 inactivation also reduces the amplitude of both pulse types (Figure 4H) [5]. This implies that although ps1 is not strictly necessary for P_fast_ production - ∼10% of pulses are still of the P_fast_ type and these P_fast_ pulses are indistinguishable from wild type P_fast_ pulses (Figure 4I, Supplemental Figure 4E, F) - this motor neuron contributes instead to the choice between pulse types, and thereby the overall structure of male song.

## Discussion

A central aim of systems neuroscience is to combine neural activation, silencing, and recording techniques to causally link particular neurons and circuits to behavior and to show how activity patterns within these neurons relate to behavioral activity. New computational methods (e.g. [12,29,30]) are facilitating the automated detection and classification of behaviors to drive these studies forward, particularly in genetic model systems, such as worms, flies and mice [31,32]. But solving the neural basis for behavior requires the appropriate temporal and spatial parsing of behaviors in order to generate meaningful connections with the underlying neural and muscle activity patterns that drive them. Here we use a highly quantifiable and robust behavior - the courtship song of *Drosophila melanogaster* - to investigate this fundamental issue. We show here using new unsupervised classification methods that song comprises at least three, not just two, modes, and we show how this distinction affects our interpretation of the mechanisms underlying song production (Figure 3, 4), perception (Figure 3J) and evolution (Figure 2G-J).

Rather than being a reflex-like behavior, song production relies on the integration of multiple sensory cues [5,9] and discriminating between P_slow_ and P_fast_ now reveals a novel layer of song control. Previous studies had shown that, similar to humans, *Drosophila* males increase the amplitude of their acoustic communication signals based on a visual estimate of distance to the receiver [5]. Analyzing the effect of distance on both pulse types separately, we now find that visual information mediates this amplitude modulation using two separate control mechanisms: First, a digital/binary control system biases pulse production towards the louder P_fast_ when far from the female (Figure 3D). And second, an analog control system upregulates P_fast_ - but not P_slow_ - amplitude with distance (Figure 3G). The strong interaction between the carrier frequency of each song mode and its amplitude (Figure 3F) suggests that mechanical constraints may facilitate the production of loud signals at higher carrier frequencies. These two modes of control are likely implemented through distinct circuits, and hence discriminating the two pulse types is important for understanding the neural basis of this behavior.

Cataloguing all song modes is also important for dissecting the neural circuits driving song production. Female proximity cues activate the song command neuron P1 [17,19] and using optogenetics, we have shown that activation of P1 biases solitary males towards singing more sine song (Figure 4B_1,2_, C_1,2_) - which is naturally produced when the male is closest to the female [9]. However, P1 activation had no strong effect on the choice between P_slow_ and P_fast_, suggesting that pulse choice is implemented elsewhere in the circuit (Figure 4B_1,2_). By contrast, pIP10 is thought to be downstream of P1 [18], but its activation strongly biases males toward producing P_fast_ instead (Figure 4B3, C3). Thus, rather than being a relay of P1 activation, pIP10 also actively shapes the dynamics of song mode choice (Figure 4B, Supplemental Figure 4A), but not the parameters of each song mode (Supplemental Figure 4C, D). This is an important distinction - these neurons appear to bias the output of the downstream pattern generating circuits without changing their control over the features of each song mode, such as the frequency or shape of individual pulses. Moreover, activation of P1 or pIP10 in solitary males produces all 3 song modes, but with different probabilities and latencies. This suggests that a static analysis of song production based on the activation or inactivation of song command neurons is insufficient to understand how song is produced in *Drosophila*. Rather, the activity dynamics of different elements of the song pathway shape what is sung and when.

ps1 was a motor neuron previously implicated in setting the frequency and amplitude of pulses [8]. By segmenting the behavior to identify multiple pulse types, we find instead that this motor neuron does not strongly affect the parameters of each song mode (Figure 4I) (similar to the results with P1 and pIP10 activation, Supplemental Figure 4C, D), but rather the choice of which pulse type to produce relative to the distance to the female (Figure 4F). This suggests that this neuron represents one of the ends of the pathway that connects visual information with pulse choice. Further studies into the activity of command neurons, motor neurons, and muscles [8,18,33,34] during singing will shed additional light onto the circuits choosing and shaping the three song modes.

We also found that females slow differentially to P_slow_ and P_fast_ (Figure 3J, Supplemental Figure 3), and that this slowing depends on the context: females reduce their locomotor speed to P_fas_t only when it is sung at farther distances (Figure 3J). This demonstrates that distinguishing between pulse types matters for studying the perception of song as well. Future experiments that allow precise control over the sensory cues available to the female - e.g. through sound playback - are necessary to elucidate the sensory source of the differential slowing response and ultimately its neural basis.

While previous studies demonstrated the existence of multiple pulse types in Drosophila species [10,35-37] our statistical analysis of pulse shapes in *D. melanogaster, D. simulans* and *D. mauritiana* now shows that P_fast_ and P_slow_ shapes are conserved in these three species (Figure 2H-J, Supplemental Figure 2), implying that the existence of these two pulse types predates the species split. Interestingly, *D. yakuba* - a member of the melanogaster group outside of the branch considered here (Figure 2G) - produces two pulse types termed “thud” and “clack” that are claimed to be produced at different distances to the female {Demetriades:1999iw}, just as with P_slow_ and P_fast_ in *D. melanogaster*. It is conceivable that the thud and clack pulses in *D. yakuba* are homologous to P_slow_ and P_fast_ but further studies are required. It is also not yet clear whether distance to the female is the main driver of pulse choice in *D. simulans* and *D. mauritiana*.

In summary, we show that segmenting Drosophila melanogaster song into three, not two, modes affects our interpretation of the sensory, neural, and evolutionary effects on song patterning. This study highlights that a detailed characterization of behavior is required to correctly interpret data from neural activation and silencing experiments, recordings of neural activity, and comparisons of behavior across species.

## Author Contributions

C, PC, and MM designed the study. JC, PC, FR, DM, TP, and DP collected data. JC, PC, FR, and TP analyzed data. JC and MM wrote the manuscript.

## Acknowledgements

We thank Asif Ghazanfar, Josh Shaevitz, and David Stern for comments on the manuscript, and Georgia Guan for technical assistance. JC was supported by a postdoctoral fellowship through the Princeton Sloan-Swartz Center, and MM was supported by an NSF BRAIN EAGER award, an NIH New Innovator Award, a McKnight Scholar Award, and a Faculty Scholar Award from the Howard Hughes Medical Institute.

## Contact for Reagents and Resource Sharing

Further information and requests for resources and reagents should be directed to and will be fulfilled by the Lead Contact, Mala Murthy.

## Experimental Model and Subject Details

### Animals

If not stated otherwise, virgin male and female flies were isolated within 6 hours of eclosion and aged for 3-7 days prior to experiments. Flies were raised on a 12:12 dark:light cycle, at 25°C and 60% humidity.

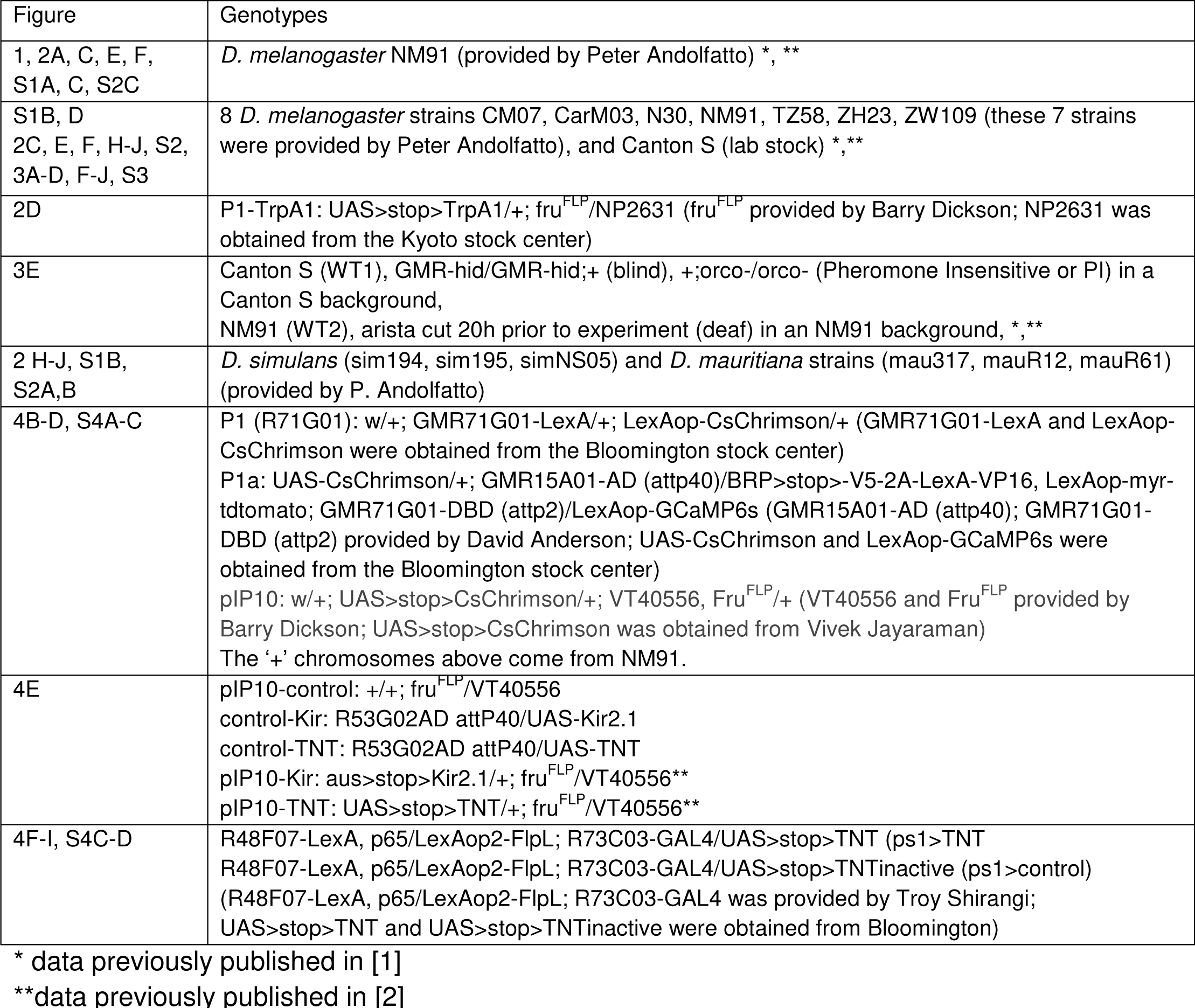

## Method Details

### Behavioral chamber

Behavioral chambers were constructed as previously described [1-3]. For optogenetic activation experiments we used a modified chamber whose floor was lined with white plastic mesh and equipped with 16 recording microphones. To prevent the LED light from interfering with the video recording and tracking, we used a short-pass filter (Thorlabs FESH0550, cut-off wavelength: 550 nm). For the analysis of fly pose during the production of all three song modes we recorded video at 100 frames per second (instead of 60 frames per second as in all other recordings) and designed a novel chamber with a domed lid and a flat floor that was lined with fine nylon mesh (Ted Pella Nylon 300 Mesh) instead of the coarse mesh used for the other recordings. This created a clean background and allowed us to resolve fine details of the fly body like the wing. When recording with a female, males were painted on the thorax with a white dot 20h prior to experiment under cold anesthesia for identification of sex during tracking. Flies were introduced gently into the chamber using an aspirator. Recordings were timed to be within 150 minutes of the behavioral incubator lights switching on to catch the morning activity peak. Recordings were stopped after 30 minutes or earlier if copulation occurred. If males did not sing in the first 5 minutes of the recording, the experiment was discarded.

### Inactivation of song command and motor neurons

Tetanus toxin (TNT) was used for chronic inactivation of the motorneuon driving the direct wing muscle ps1 [4], with an inactivated TNT (TNT-in) as a negative control. Tetanus toxin (TNT) and the inward-rectifying Potassium channel K_ir_ 2.1 were used to inactivate the song command neuron pIP10 during courtship [5].

### Optogenetic activation

Flies were kept for at least 3 days prior to the experiment on fly food supplanted with retinal (1ml all-trans retinal solution (100mM in 95% ethanol) per 100ml food). CsChrimson [6] was activated using a 627 nm LED (Luxeon Star) at an intensity of 0.46 mW/mm^2^ (driving voltage 3.2V). Stimulus period 3sec, 9 duty cycles (DCs) ranging between 0.1 and 0.9, filling between 300 and 2700ms of the period with constant LED illumination. Each DC was presented in randomized blocks of 90s (30 trials each). Since optogenetic activation experiments were performed using solitary males, recordings were not timed to peaks in circadian activity. Sound recording and video were synchronized by positioning a red LED that blinked with a predetermined temporal pattern in the field-of-view of the camera and whose driving voltage was recorded alongside the song.

## Quantification and Statistical Analysis

### Song segmentation and tracking

Both tracking of flies and segmentation of song recordings were performed as previously described [1-3]. Pulses were detected with high detection rates (true positive rates 78%, 74%, 85% for *D. melanogaster, simulans* and *mauritiana)* and few false detections (false positive rate ∼2% for *D. melanogaster* and *D. simulans*, 6% for *D. mauritiana* because some pairs produced substantial movement noise during courtship) (Supplemental Figure 1B). The song segmenter (with the parameters used in [1], not those of [7] - see [8] for a comparison of segmenter performance with these different parameters) for *D. melanogaster* detected P_fast_ and P_slow_ with similar high rates without requiring any modifications (77% and 80%, respectively). Note that we extended the song segmenter to now also classify pulses into P_fast_ and P_slow_ (available at https://github.com/murthylab/songSegmenter, see below). For segmenting the pulses produced by *D. simulans* and *D. mauritiana* we modified the segmenter by building strain specific pulse models from sets of manually annotated data (∼500 pulses per strain).

### Waveform selection for segmented pulses

For *D. melanogaster*, pulse waveforms were extracted from the recordings by taking 25.1 ms (251 samples at 10kHz) around the pulse center detected by the song segmenter from the channel on which the pulse was recorded with the highest energy. For the much shorter pulses produced by *D. simulans* and *D. mauritiana*, 25.1 ms around the pulse peak contained mostly noise. We therefore upsampled the waveforms to 20kHz and extracted the same 251 samples - now corresponding to 12.55 ms - around the pulse center.

### Waveform selection for unbiased analysis of song waveforms

For the unbiased analysis of song structure, we selected waveforms from the raw recordings without using the song segmenter. As a first step, all 9 recording channels were merged by choosing the signal from the channel with the highest absolute value in 5ms windows. To avoid our analysis being drowned out by background noise (up to 90% of a song recording can contain no song), we chose only signals that exceeded the energy of the recording noise. Sound energy was calculated by estimating the signal envelope using the Hilbert transform, transforming it to a logarithmic scale and then smoothing it using a sliding window lasting 100ms. The logarithmic transformation produced an energy distribution that resembled a mixture of two Gaussians, one corresponding to baseline noise and one corresponding to song signals. We fitted a Gaussian mixture model with two components to the distribution energy values and identified as “signal” the component with the higher mean energy. From all time points classified as signal we then extracted 25ms long, non-overlapping waveforms. Note that the resulting data set contained many snippets that contained only noise if they were at the very beginning of or between song bouts.

### Waveform normalization

To remove variability in the waveforms arising from the position and distance of the singing male from the microphone, we

1. **Divide** the raw waveforms, *x(t)*, by their norm: *x(t)/√∑x(t)*^2^.
2. **Center** to their peak energy obtained by smoothing the square pulse waveform with a rectangular window spanning 15 samples (1.50 ms for *D. melanogaster*, 0. 75 ms for *D. simulans* and *D. mauritiana*).
3. **Flip** sign such that average pulse in the 10 samples (1.00 ms for *D. melanogaster*, 0.5 ms for *D. simulans* and *D. mauritiana)* preceding the pulse center is positive.

Time scales were shortened for *D. simulans* and *D. mauritiana* to enable robust alignment of the much shorter pulses produced by these species (see e.g. Figure 2H). The last step - adjusting waveform sign - does not artefactually create clustering into two pulse types but rather reduces the degeneracy of the waveform space since the same pulse can occur with different signs on different microphones, indicating that the position of the male wing relative the microphone (in front of or behind) determines waveform sign. Clustering waveforms with randomized sign yields clustering in which each pulse type appears in two clusters - one in its original shape and one with the sign inverted (Supplemental Figure 2C).

### Dimensionality reduction

The dimensionality of the normalized waveforms was reduced from 251 data points per waveform to two dimensions using two different methods:

1. **T-distributed stochastic neighbor embedding (tSNE)** preserves local neighborhood structure while ignoring global similarity between data points. It is therefore ideally suited for cluster analysis, which relies on local similarity. To embed large data sets (more than 100000 pulses) we chose the computationally and memory-efficient Barnes-Hut implementation [9] (parameters: initial PCA dimensions=30, perplexity=50, accuracy=0.1). The shape of the embedded distribution was highly reproducible across runs and did not depend critically on the choice of parameter values. Since large perplexity values increased the separation between clusters - while preserving the overall shape of the embedded distribution - we chose a perplexity value of 50.
2. **Principal component analysis (PCA)** is a linear method for dimensionality reduction and works by finding a small set of orthogonal basis functions in which the global data variance is maximized. Clustering using a PCA-based representation of the waveforms yields qualitatively similar results (Figure 2C), indicating that our findings do not critically depend on the nonlinear tSNE embedding. However, the cluster separation corresponding to P_slo_w and P_fast_ was not as strong and hence clustering was not as reproducible as with tSNE. Consequently, we chose to use tSNE embedding for all analyses.

### Waveform clustering

To identify groups of similar pulses, we clustered the waveforms in two steps:

1. The 2D distribution of data was partitioned using the watershed algorithm, which cuts a distribution along local minima. To that end we estimated the distribution of the tSNE-embedded pulses using kernel density estimation (matlab function mvksdensity), with the bandwidth chosen automatically by the algorithm (Figure 1C). This resulted in seven clusters (thin lines in Figure 1C and cluster waveforms in Figure 1D), some of which exhibited highly similar mean waveforms (Figure 1E) indicating that this first step overpartitioned the waveform space.
2. We therefore consolidated these watershed clusters using hierarchical clustering of the cluster centroids (matlab function clusterdata). When clustering only the pulses detected by the song segmenter (Figure 2A), the resulting cluster trees always revealed a simple bipartite structure and we hence chose a clustering cutoff that produced two modes. For the unbiased analysis of song waveforms (Figure 1D), the cluster tree revealed four principal waveform shapes corresponding to noise, sine song and the two pulse types. Cluster consolidation

a. joins three sine clusters, which is supported by them containing all similar waveforms (Figure 1E, left)
b. joins two P_slow_ clusters but leaves the P_fast_ separate, which is again supported by the waveforms in the P_slo_w clusters being highly similar and those in the P_fast_ cluster being highly dissimilar to those of P_slow_.

Note that there exists a myriad of approaches for clustering data. Watershed plus consolidation using hierarchical clustering on the centroids produced the most robust clustering. Direct agglomerative clustering of the tSNE embedded data - without first using the watershed cluster step - also results in four clusters (not shown), but these clusters are not as “clean” - for instance, the P_fast_ mode extends into the noise cluster. Direct clustering on the unbiased set of waveforms yields unreliable results, likely because of the large variability of waveforms in this data set (it includes noise).

The full analysis pipeline yielded near-identical results across different runs - individual pulses were almost always assigned to the same cluster with very high probability (adjusted Rand index ∼0.97). Code illustrating all steps of the pipeline is available at https://murthylab.github.io/pulseTypePipeline/.

### Template-based pulse type classifier

Normalized pulses from the *D. melanogaster*wild type strain NM91 were projected onto the centroids of the P_slow_ and P_fast_ clusters and a quadratic boundary separating both pulse types was obtained using quadratic discriminant analysis cross-validated using an 80:20 partition into training and test data (Supplemental Figure 1C). The classifier performs well on the test data and generalizes well to different *D. melanogaster* strains (Supplemental Figure 1D). It is therefore a fast and robust method for classifying pulses that does not require the computationally extensive pipeline outlined above. Code for building and evaluating the classifier is available at:

https://github.com/murthylab/pulseTypeClassifier. The classifier has also been integrated into the song segmenter software (available at https://github.com/murthylab/songSegmenter).

### Characterization of pulse shapes

Song pulses are transient signals and their raw magnitude spectra are relatively broad (Figure 2B, Supplemental Figure 4E). Accordingly, the *peak* frequency of the spectrum is an unreliable measure of pulse carrier frequency. We hence used the center of mass of the magnitude spectra thresholded at e^-1^=0.37 as a measure of frequency. The values obtained tended to closely resemble 1) the spectral peak frequencies in cases of sufficiently peaked spectra (e.g. for most instances of P_slow_) and 2) the carrier frequency values extracted from fits of Gabor functions to the pulse waveforms (not shown), demonstrating that the center of mass faithfully extracts the pulse carrier frequency. A pulse symmetry index was calculated as the dot product of the first half of the pulse and the flipped second half of the pulse. Positive indices correspond to even symmetry and negative indices to odd symmetry. Pulse amplitudes were normalized for the gain of individual microphones as described in [2].

### Identification of sensory cues driving pulse choice

The GLM analysis for identifying the song features driving the choice between P_slow_ and P_fast_ was performed as in [2].

### Amplitude modulation plots

We evaluated the relation between pulse amplitude and distance to the pulse at a delay of 470ms since the distance at this delay is most predictive of pulse amplitude [2]. Distance was binned between 1.75 and 10.25 mm in 0.5mm steps and the mean pulse amplitude (and its standard error) were calculated for each.

The first three pulses of each train were excluded from the analysis since flies do not modulate pulse amplitude at pulse train start [2]. Distance bins with <50 pulses across all flies were excluded before calculating r^2^ values.

### Analysis of optogenetic activation data

The song probability traces (Figure 4B, Supplemental Figure 4A) were constructed by calculating the fraction of trials (for each time bin, pooled across all fies) during which a male produced sine, P_slow_ or P_fast_. Traces do not add up to 1.0 since there are trials during which males did not sing any song. Song probability traces and male speed were down sampled to 400Hz and smoothed with a sliding Gaussian kernel lasting 150ms with standard deviation σ=64ms.

## Data and Software Availability

Song segmenter with pulse classifier: https://github.com/murthylab/songSegmenter.

Stand-alone pulse classifier: https://github.com/murthylab/pulseTypeClassifier.

Script illustrating the analysis pipeline: https://murthylab.github.io/pulseTypePipeline/.

Raw data is available upon reasonable request from the Lead Contact, Mala Murthy

## Supplemental Figures

**Supplemental Figure 1:**
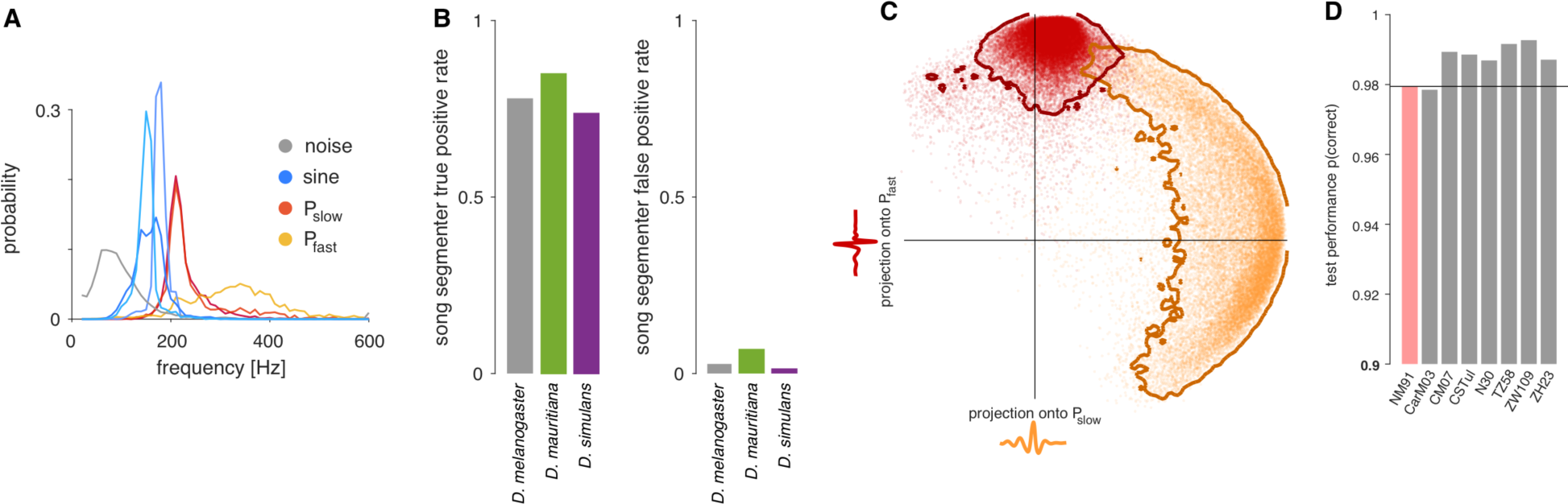
P_fast_ and P_slow_ are distinct song modes. **A** Distribution of carrier frequencies for all 21,104 waveform shapes in Fig. 1C, separated by watershed clusters from Fig. 1E (see legend for color code). **B** True (left) and false (right) positive rates for the automatic segmentation of pulses from ***D. melanogaster*** (grey), ***D. mauritiana*** (green), and ***D. simulans*** (purple). True positive rates (left) are high for all species (>74%) with only few false positives (right, <7%). ***D. mauritiana*** (green, left) exhibits slightly elevated false positive rates because this species tends to produce a lot of movement noise during courtship. **C** A template-based pulse type classifier was trained by projecting each pulse onto the P_slow_ and P_fast_ centroids obtained from the cluster analysis. Each dot represents on pulse colored by the pulse label assigned in the cluster analysis. Red and orange lines correspond to the contour enveloping 95% of the pulses for each pulse type. A quadratic decision boundary is then determined using Quadratic Discriminant Analysis (QDA) to classify novel pulses based on their projection values. **D** Test performance of the QDA classifier for the strain on which the classifier was trained on (NM91, red) - using a set of pulses not used for training - and for 7 completely new ***D. melanogaster*** wild type strains (grey). Note the y-axis starts at 0.9 to highlight differences between strains. The classifier generalizes excellently across wild type strains of ***D. melanogaster***.

**Supplemental Figure 2:**
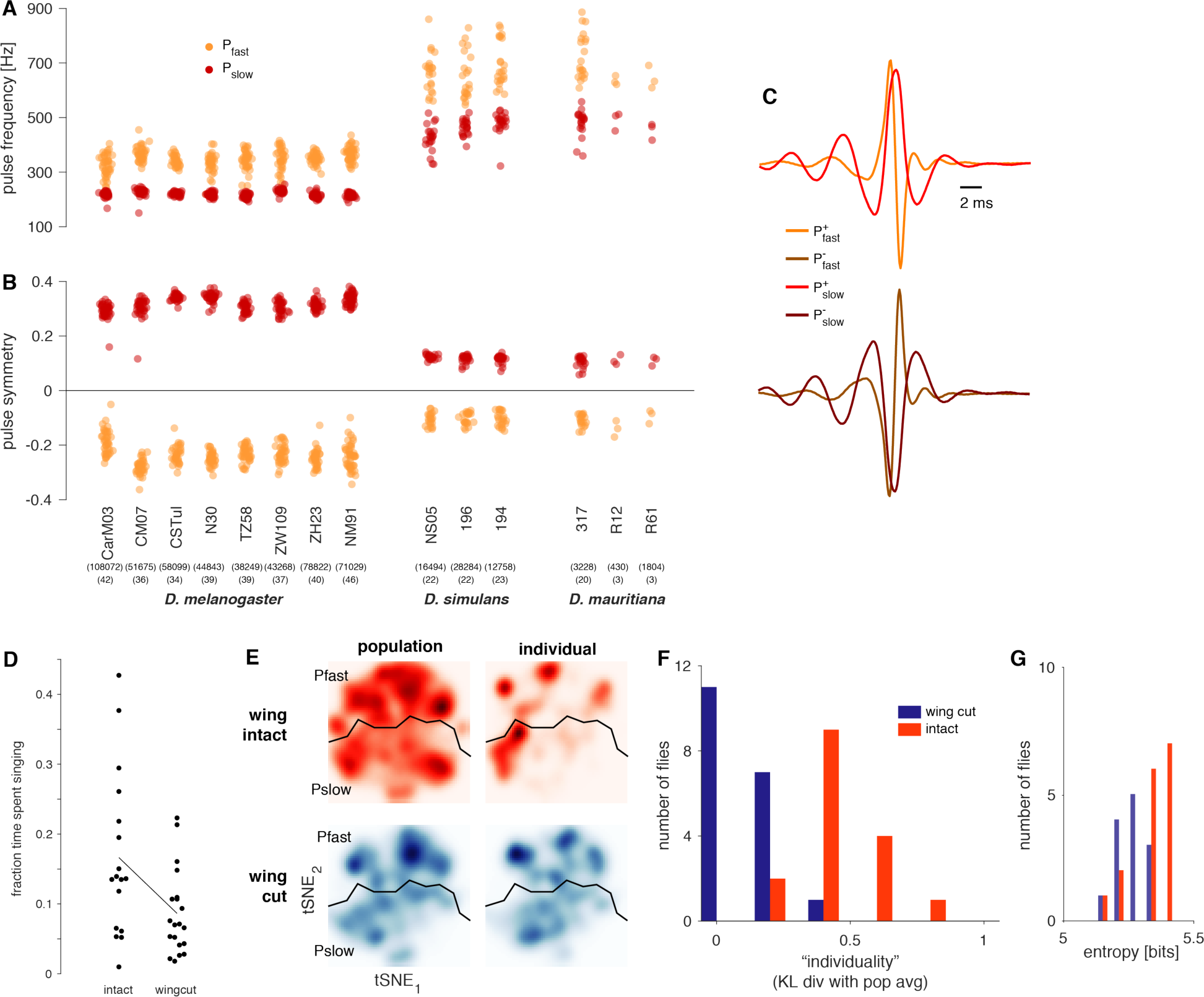
P_fast_ and P_slow_ are produced by strains of several *Drosophila* species. **A, B** Pulse carrier frequency (A) and pulse symmetry (B) for P_fast_ (orange) and P_slow_ (red) centroids of all individuals for the 8 *D. melanogaster*, 3 *D. simulans*, and 3 *D. mauritiana* strains. P_slow_ tends to be the slow and symmetrical, and P_fast_ tends to be the fast and asymmetrical pulse type in all individuals. Numbers in parentheses indicate the number of pulses (top) and flies (bottom) tested for each strain and species. **C** Clustering pulses with randomized sign produces clusters in which each pulse type appears in a version with “correct” (top, bright colors) and inverted (bottom, darker colors) sign. **D** Wing cut males (N=20 pairs, a single wing of the male (either left or right) cut 24h before the experiment) produce less song than intact controls (N=17 pairs) (p=0.01, two-tailed Wilcoxon rank sum test). **E** All pulses (wing intact - red, top; single wing cut - bottom, blue) were embedded into the same tSNE space and clustered to identify P_slow_ and P_fast_ pulses (boundary between pulse types is marked by a thick black line). We then compared the distribution of pulses in tSNE space between the population (left) and individuals (right) (distributions where estimated as for the pulse clustering, see STAR Methods for details). Wing cut males (bottom) produce pulses with a distribution that tends to resemble that of the population. By contrast, the pulses of intact males seem to be more “idiosyncratic” than those of wing cut males - meaning the pulse distribution in intact males is very dissimilar to that of the population. **F** We quantified the similarity of the pulse distributions of individuals and the population in tSNE space using the Kullback-Leibler divergence (KL) for both pulse types: DKL(PPOP, pind) = Ippoplog2(ppop/pind) where ppop and pind are the distributions of pulses in tSNE space for the population and for individuals, respectively, as shown in E. Wing cut flies (blue bars) have smaller KL values (meaning higher similarity to the distribution of the population) than intact flies (red bars) (p=2×10^-5^, two-tailed Wilcoxon rank sum test). **G** The variability of pulse shapes - given by the entropy of the distribution of pulse shapes in tSNE space - changes only little, indicating that the lack of a wing does not strongly affect the diversity of pulses produced (p=6x10^-6^, two-tailed Wilcoxon rank sum test).

**Supplemental Figure 3:**
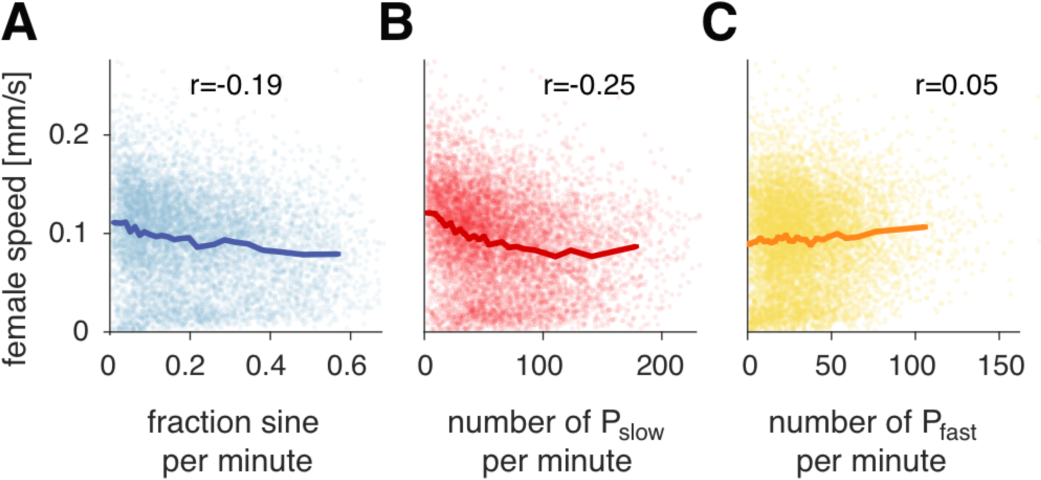
Correlation between song modes and female speed. **A, B, C** Rank correlation between female speed and amount of sine (A) or number of Psiow (B) or P_fast_ (C) pulses per 30 second time window of courtship (8826 windows from 315 *D. melanogaster* pairs used). If receptive, females slow down in response to song {Coen et al. 2014}. Individual points correspond to individual time windows, shaded lines are the binned averages (mean ± SEM). Bin width was chosen adaptively such that each bin contains the same number of data points. Only the amount of sine song and P_slow_ are strongly correlated with a reduction in female speed (rank correlation values indicated in each panel, sine r=-0.19, p=2×10^-74^; P_slow_ r=-0.25, p=2×10^-129^; P_fast_ r=0.05, p=2×10^-5^).

**Supplemental Figure 4:**
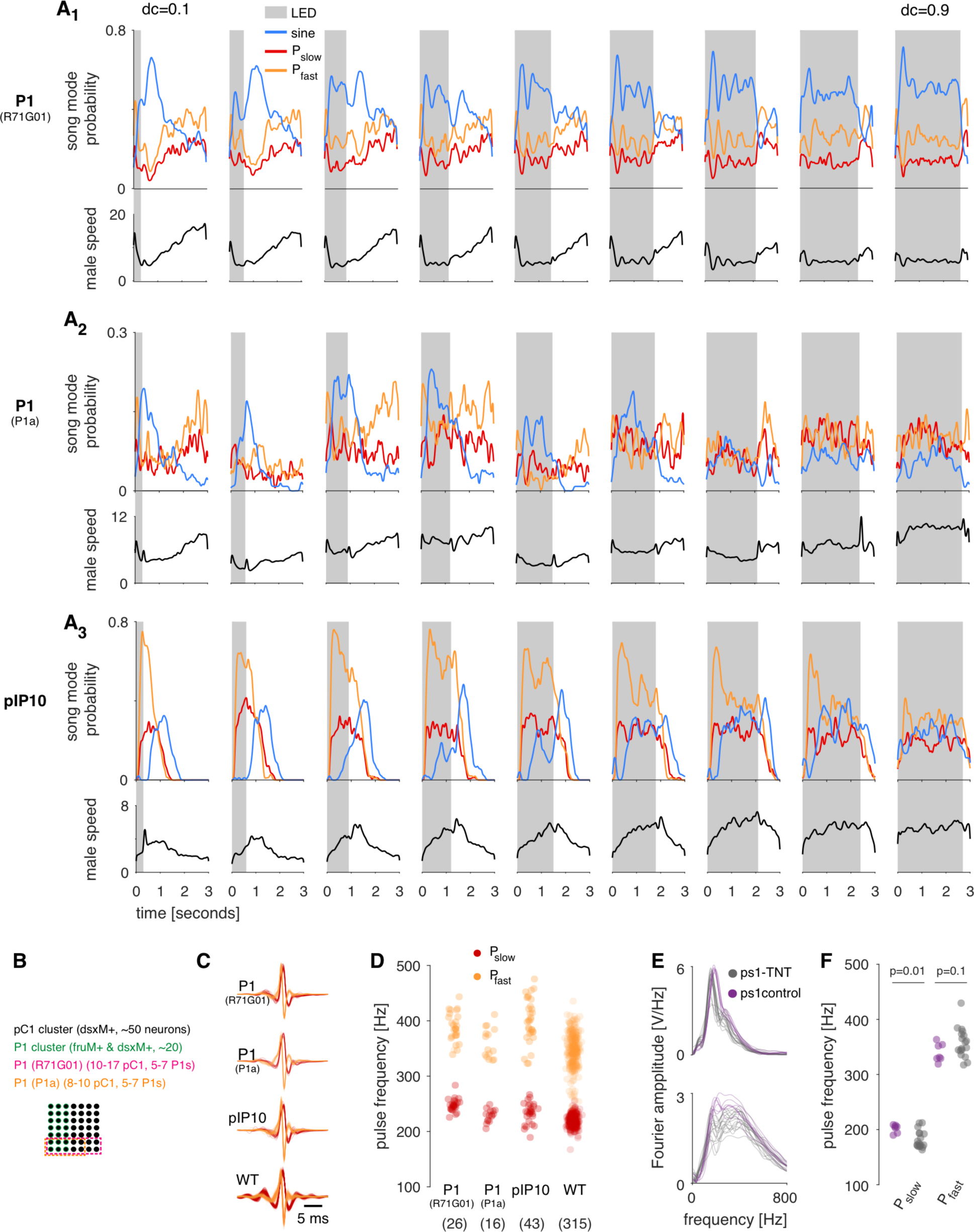
Three different song pathway neurons and their role in song patterning. **A** Population average song probability for producing sine song (blue), Psiow (red) or ft (orange) upon optogenetic activation of P1 (Ai), P1a (A2), and pIP10 (A3). Same data as in Fig. 1B in the main manuscript but with all nine DCs used. Black lines show the trial and population averaged male speed. **B** Schematic of the labelling of P1 (R71G01) and P1 (P1a). pC1 (black) is an anatomical cluster of neurons that are *DsxM+*. P1 (green) is a subset of pC1 that is also *FruM+*. R71G01 [5] (labelled “P1 (R71G01)” in our figures) labels 5-7 P1 neurons, in addition to 10-17 pC1 neurons and many other neurons in the fly. P1a [6] labels the same set of P1 neurons as R71G01 but only 8-10 FruM-pC1 neuron and a few other neurons. **C, D** P_slow_ (red) and P_fast_ (orange) shapes (C) and frequencies (D) for optogenetic activation of P1 (R71G01), P1a, and pIP10. *D. melanogaster*wild type data shown for comparison. Number of flies for panels C and D are given in parentheses in panel D. **E** Spectra of P_slow_ (top) and P_fast_ (bottom) in ps1-ctrl (purple) and ps1-TNT flies (grey). Note the small peak shift for P_slow_ for ps1 inactivation. **F** Frequency of P_slow_ (right) and P_fast_ (left) pulses for ps1 control (purple) and ps1 experimental (gray) flies. Each dot corresponds to the frequency of the average pulse for one individual (N=24 experimental flies (36358 pulses) and 8 control flies (25850 pulses)). Inactivation of ps1 decreases P_slow_ but not P_fast_ frequency (P_fast_, p=0.1; P_slow_ p=0.01, two-tailed Wilcoxon rank sum test). E-F: N=7 experimental flies (36358 pulses) and 24 control flies (25850 pulses).

